# Sustained Release of IL-2 Using an Injectable Hydrogel Prevents Autoimmune Diabetes

**DOI:** 10.1101/2020.03.15.993063

**Authors:** Nadine Nagy, Gernot Kaber, Michael J. Kratochvil, Hedwich F. Kuipers, Shannon M. Ruppert, Koshika Yadava, Jason Yang, Sarah C. Heilshorn, S. Alice Long, Alberto Pugliese, Paul L. Bollyky

**Author notes:** Corresponding author: Nadine Nagy, PhD, Department of Medicine, Division of Infectious Diseases, Stanford University, 279 Campus Drive, Beckman Center, Room B237, Stanford CA 94305, USA.

## Abstract

Interleukin 2 (IL-2) is a promising therapy for autoimmune type 1 diabetes (T1D), but the short half-life *in vivo* (less than 6 minutes) limits effective tissue exposure to IL-2. Tissue exposure is required for the tolerogenic effects of IL-2. We have developed an injectable hydrogel that incorporates heparin polymers to enable the sustained release of IL-2. This platform uses clinical grade and commercially available materials, including collagen, hyaluronan, and heparin, to deliver IL-2 by slowly degrading and releasing IL-2 over a two-week period *in vivo*. We find that heparin potentiates the activity of IL-2 and IL-2-mediated expansion of Foxp3+ regulatory T cell (Treg). Hydrogel-mediated IL-2 release showed a reduction of CD4+ and CD8+ T cells and an increase of FoxP3+ Treg in the lymph nodes of injected mice. Moreover, in the Non-Obese Diabetic (NOD) mouse model of T1D once-weekly administration of IL-2 hydrogels prevented diabetes onset as efficiently as 3x weekly repeated injections of soluble IL-2. Together these data suggest that heparin-containing hydrogels may have benefit in delivering low-dose IL-2 and promoting immune tolerance in autoimmune diabetes.

## INTRODUCTION

Type 1 diabetes (T1D) is characterized by the progressive immune cell-mediated destruction of pancreatic β-cells and the failure of regulatory mechanisms that normally prevent this, including Foxp3+ regulatory T cells (Tregs) (1-3). The absence or depletion of Treg leads to multi-systemic autoimmunity (1) whereas their adoptive transfer prevents autoimmunity (4).

One critical factor that governs Treg function is the cytokine interleukin 2 (IL-2) (5,6). Low-dose IL-2 injections are a promising therapeutic approach in T1D and other autoimmune diseases (7,8). However, IL-2 has a short half-life and requires daily dosing (9). It would be desirable to administer IL-2 in a sustained manner with as few injections as possible, and localize the administration of IL-2 to reduce off-target effects, including eosinophilia and activation of CD56hi NK cells.

One matrix component that has been used to deliver cytokines and growth factors in a sustained manner is heparin (HI), a sulfated glycosaminoglycan. HI and its analog heparan sulfate bind a diverse range of cytokines, chemokines and growth factors (10,11). HI-containing hydrogels containing various factors have been developed for stem cell culture and other applications (12,13). HI-binding also influences the bioactivity of these molecules (14,15). HI is used at low concentrations in these formulations (typically <0.1% weight/volume), meaning that the anti-coagulant properties of HI are not an issue. Beyond localized delivery of cytokines, HI enhances the bioactivity of IL-2 (16,17) though effects on Treg have not been previously examined.

We hypothesized that it would be possible to adapt a HI-based hydrogel for the sustained release of IL-2 to prevent autoimmune diabetes in the non-obese diabetic (NOD) mouse model of T1D.

## RESEARCH DESIGN AND METHODS

### Mice

All animals were bred and maintained under specific pathogen-free conditions, with free access to food and water, in the vivarium at Stanford University (Stanford, CA). Female NOD and C57Bl6 mice were purchased from The Jackson Laboratory (Bar Harbor, ME). All experiments and animal use procedures were approved by the Animal Care & Use Committee at Stanford University.

### Weight and diabetes monitoring

Beginning at 4 weeks of age, mice were weighed and bled weekly to measure their blood glucose levels (Contour, Bayer Healthcare, Tarrytown, NY). When two consecutive blood glucose readings of 250 mg/dL or greater were recorded, mice were considered diabetic.

### Hydrogels

Extracel and Extracel HP hydrogels (BioTime, Inc; Alameda, CA) were generated as per the manufacturer’s instructions to form 1% Collagen (COL)/hyaluronan (HA)/heparin (HI) gels or COL/HA gels. To assess the stability of these *in vivo*, in some experiments hydrogels of 200 µL volume incorporating an Alexa fluor 790 fluorescent tag (Thermo Fisher, Waltham, MA) were injected SC or IP (as indicated) into mice and allowed to polymerize *in situ*. Residual hydrogel mass was then assessed at 1, 5 and 12 days post injection using an IVIS 100 *in vivo* imaging system (Perkin Elmer, Waltham, MA).

### IL-2 release assay from Hydrogels

Hydrogels were cast into a 96-well plate with 100 µL per well. After the hydrogels were formed, 200 µL of 1600 IU IL-2/mL PBS was added per well and incubated overnight. The loading solution was collected for later analysis. Release was conducted by incubating the samples in 200 µL of PBS, which was collected and replaced at the specified time points. IL-2 concentration were measured using an IL-2 ELISA (BioLegend, San Diego, CA).

### IL-2 proliferation assay

CTLL2 cells (ATCC, Manassas, VA), a T cell line exquisitely dependent on IL-2 for their proliferative capacity, were cultured for 48 hrs in the context of increasing concentrations of IL-2 with or without HI. Proliferation was measured by a Reliablue Cell Viability Reagent (ATCC, Manassas, VA).

### Isolation and analysis of leukocyte populations

T cells were isolated from spleen and lymph node cell suspensions using a CD4 isolation kit (Stemcell, Vancouver, Canada) as previously described. For T cell activation and Foxp3+ induction studies, 1×10^5^ cells were cultured with plate bound anti-CD3, soluble anti-CD28 with or without soluble IL-2 and TGFβ as indicated using previously published protocols (18). Cells were stained and analyzed via flow cytometry as previously described (19).

### Statistical analysis

Data are expressed as means +/- SEM of n independent measurements. Significance of the difference between the means of two or three groups of data was evaluated using the Mann-Whitney U test or one-way ANOVA with Bonferroni posttest, respectively. The statistical significance of differences between two groups was calculated by the Kruskal-Wallis test. A p value less than <0.05 was considered statistically significant.

## RESULTS

### HI potentiates IL-2 Treg induction

Given that HI binds IL-2, we asked whether HI-bound IL-2 potentiates the cytokine’s effects on Treg. *In vitro* we find that HI complexed with IL-2 drives proliferation more than 2-fold greater than IL-2 alone (**Figure 1A**).

**Figure 1:**
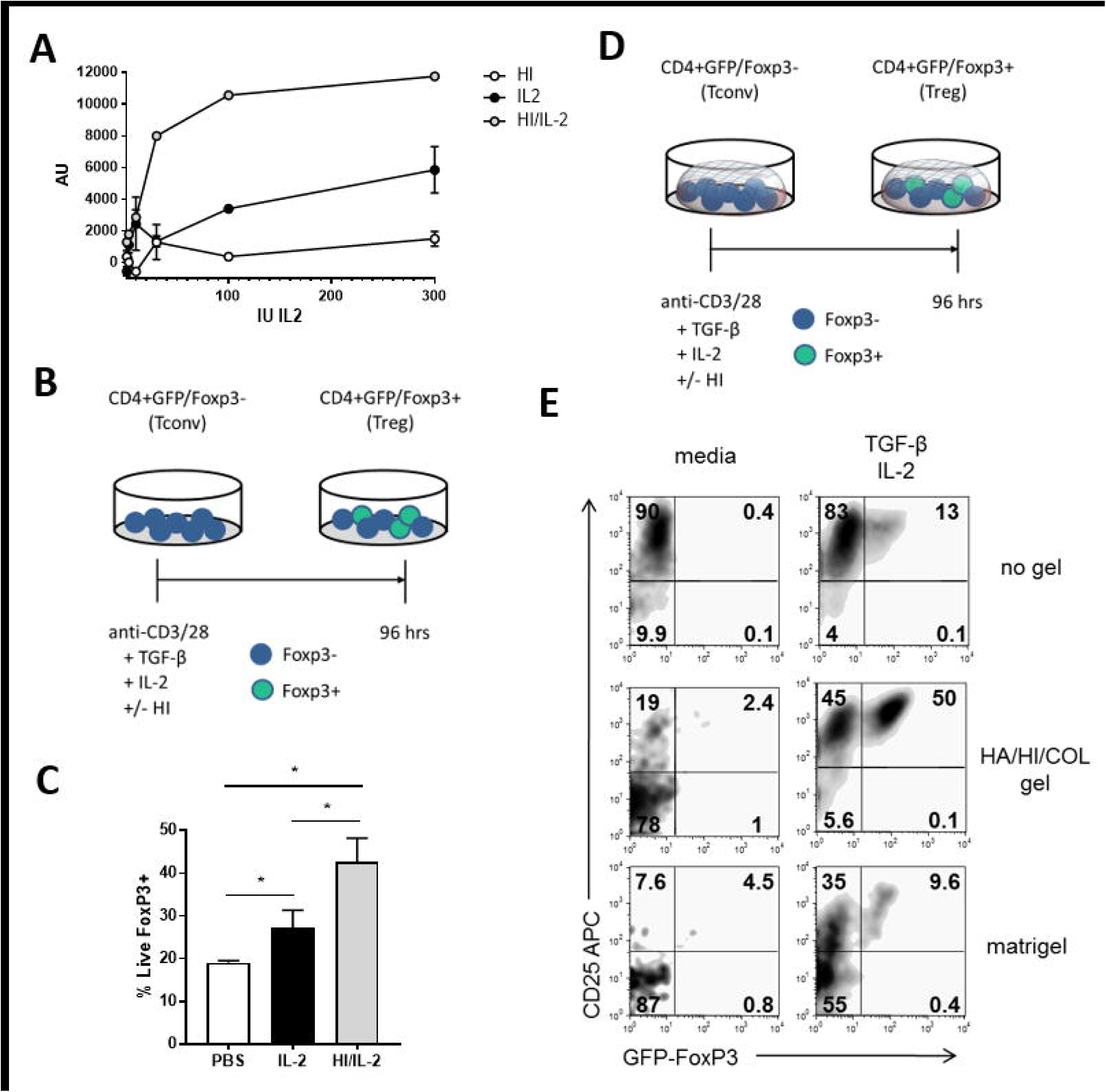
Heparin (HI) containing hydrogels potentiate Treg induction. **A.** CTLL2 cells were cultured in the context of increasing concentrations of IL-2 with or without HI. Proliferation was measured by resazurin incorporation and expressed as arbitrary units. Data for triplicate wells. **B.** Schematic of CD4+GFP/FoxP3 T cells isolated from healthy (non-diabetic) mice, cultured in the setting of anti-CD3/28, TGFβ and IL-2 in the absence or presence of HI. **C.** % Live FoxP3+ cells from experiment described in **B. D**. Schematic of CD4+GFP/FoxP3 T cells, isolated from healthy (non-diabetic) mice cultured in the setting of anti-CD3/28, TGFβ and IL-2 in the absence or presence of HI and hydrogel. **E**. % Live FoxP3+ cells measured in the different conditions as described in **D**. Representative data are shown for the mean of triplicate wells. Data represent mean +/- SEM, *P < 0.05 as determined by ANOVA.

We then sought to determine whether HI delivery likewise enhances the impact of IL-2 on Treg induction (20). CD4+GFP/Foxp3-conventional T cells (Tconv), were activated with anti-CD3/28 in the presence of TGFβ and IL-2 with or without HI (**Figure 1B**). The presence of HI and IL-2 greatly increased Treg induction in this assay (**Figure 1C**).

We next asked whether HI in the context of a hydrogel also potentiates IL-2 effects on Treg induction. We coated tissue culture plates with thin, 2-dimensional hydrogel material and performed a Treg induction assay (**Figure 1D**). We find that a HA/HI/COL containing hydrogel alone does not increase Treg induction (**Figure 1E, left column**). However, in the presence of TGFβ and IL-2 there was a significant increase in Treg (**Figure 1E, right column)**. Conversely, a matrigel and a fibrin containing gel did not potentiate Treg induction (**Figure 1E, Supplemental Figure 1**) nor did a HA/COL gel lacking the HI component (**Supplemental Figure 1**).

Together these data indicate that HI potentiates IL-2 effects on Treg expansion, and that this stimulus can be delivered in the context of a hydrogel.

### Hydrogels release IL-2 in a sustained manner and persist *in vivo*

We next sought to quantify the capacity of a hydrogel to bind and slowly release IL-2 over time. To this end, we used a commercially available COL/HA/HI hydrogel preparation (**Figure 2A**) and loaded those with 1600 IU/mL IL-2 (approximately 20,000 pg IL-2). We found that the HA/HI/COL hydrogels eluted IL-2 through day 12 while HA/COL hydrogels eluted IL-2 through day 7 (**Figure 2B,C**). These data indicate that hydrogels retain IL-2 and release it over time and that HI inclusion within the hydrogel potentiates this.

**Figure 2.**
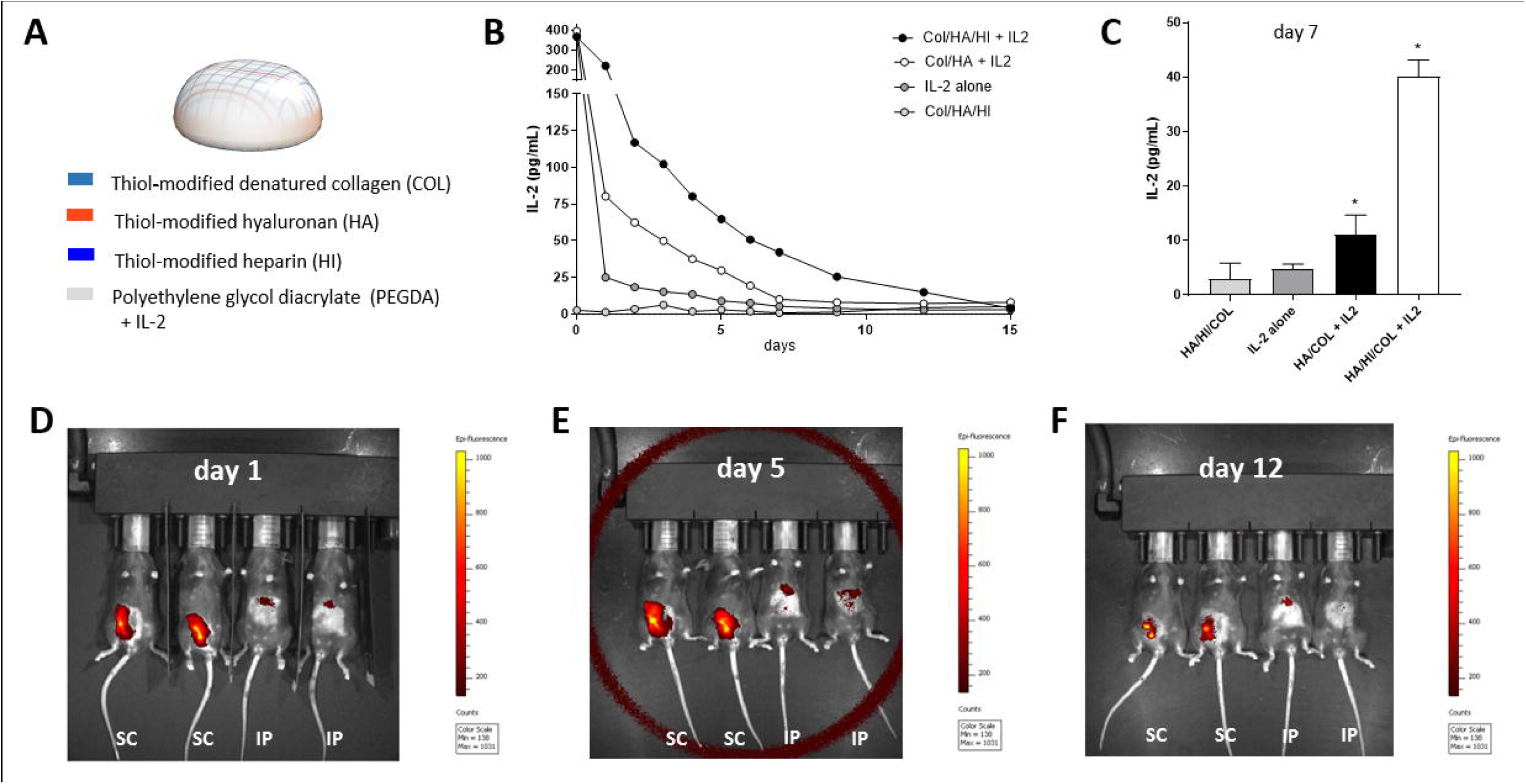
HI hydrogels release IL-2 in a sustained manner and persist *in vivo* over time. **A.** Schematic of hydrogel composition. **B.** Hydrogel IL-2 release curve for different hydrogel compositions and IL-2 alone. **C.** IL-2 release measured by IL-2 ELISA at day 7. Data are shown for the mean of triplicate wells. Data represent mean +/- SEM, *P < 0.05 as determined by ANOVA. **D-F.** Hydrogels of 50 µl volume incorporating an Alexa fluor 790 fluorescent tag were injected into mice s.c. and i.p. and allowed to polymerize *in situ*. Residual hydrogel mass was then assessed at (**D**) 1, (**E**) 5 and (**F**) 12 days post injection using an IVIS *in vivo* imaging system. Data are representative of three independent experiments.

We next asked whether injected HA/HI/COL hydrogels are stable *in vivo*. To this end, the gels were delivered as both an intraperitoneal (i.p.) and a subcutaneous (s.c.) injection and allowed to polymerize *in situ*. Residual hydrogel mass in the mice was then assessed using the IVIS *in vivo* imaging system at 1, 5, and 12 days post hydrogel injection (**Figure 2D-F**). S.c. injection of the hydrogel resulted in longer persistence, which was still clearly seen at day 12 post-injection (**Figure 2F**). These data demonstrate that a HA/HI/COL gel polymerizes *in vivo* and is stable for up to 12 days when administered s.c..

### Hydrogel-mediated IL-2 delivery is associated with a Treg increase *in vivo*

We then asked whether hydrogel-mediated IL-2 delivery could be used to promote Treg expansion *in vivo*. We chose 25,000 IU IL-2, because it was used in a previous publication to expand Treg populations and prevent progression to autoimmune diabetes in NOD mice (21). We also treated one group of mice with IL-2/IL-2 antibody complexes as a positive control. In this study NOD mice were treated with either 25,000 IU IL-2 once weekly, 25,000 IU IL-2 three times per week, or 25,000 IU IL-2 delivered in a HA/HI/COL hydrogel once weekly. Mice received this treatment regimen for one month, and T cell populations in the mesenteric lymph nodes (LN) and spleen were then assessed via flow cytometry (**Figure 3, Supplemental Figure 2).**

**Figure 3.**
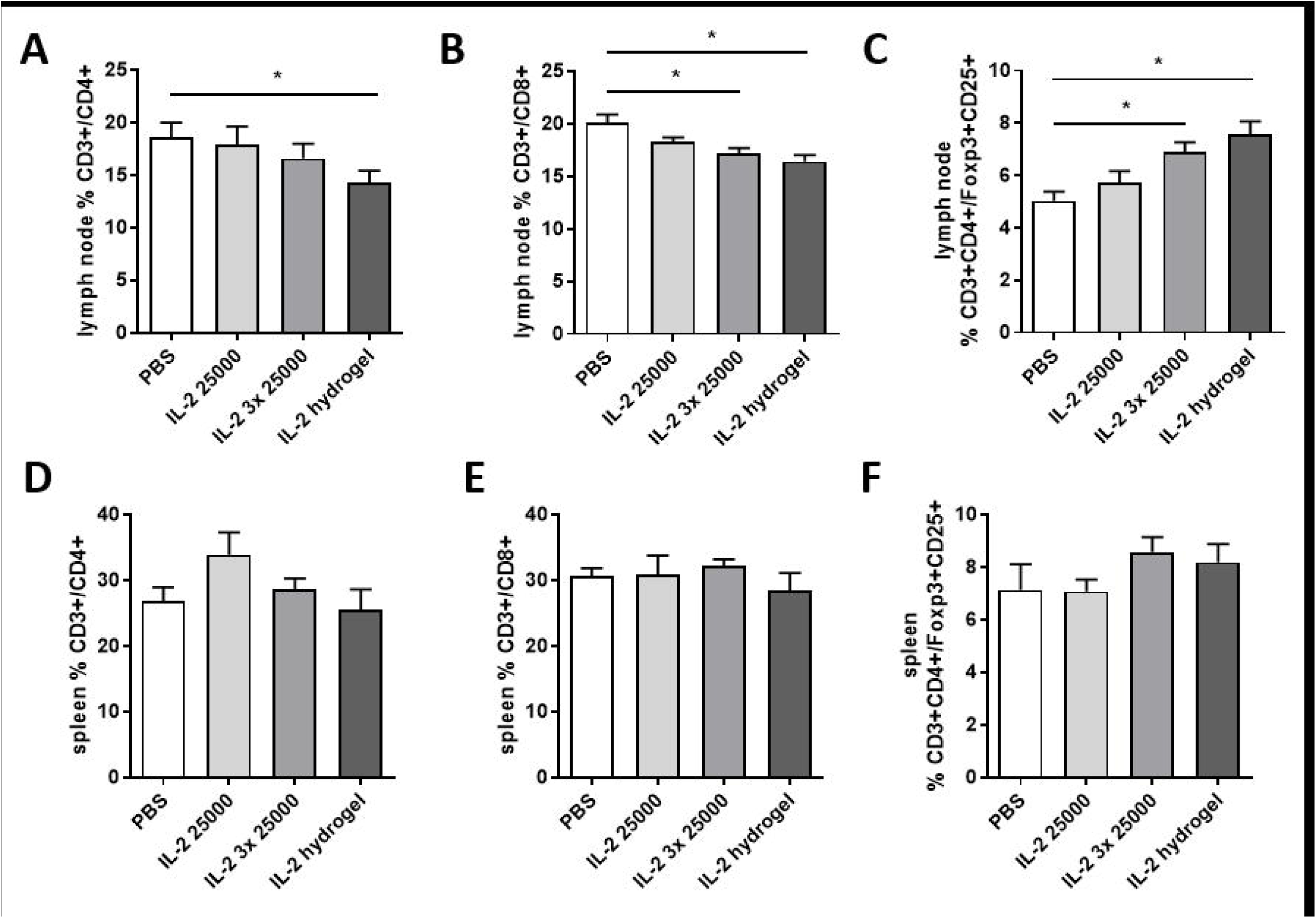
Hydrogel-mediated IL-2 delivery is associated with an increase in FoxP3+ Treg numbers. 6-week-old NOD mice were treated with either 25,000 IU IL-2 once weekly or three times a week, the same amount of IL-2 delivered in the context of a single hydrogel injection, or PBS injections as a negative control. After one month, mice were euthanized and both mesenteric lymph nodes (**A-C**) and spleens (**D-F**) were collected and populations of lymphocytes were assessed by flow cytometry. In particular, the percentage of CD3+CD4+ T cells (**A,D**), the percentage of CD3+CD8+ T cells (**B,E**) and CD3+CD4+FoxP3+ Treg (**C, F**), Data are representative of 2 independent experiments. N = 10 mice per group. Data represent mean +/- SEM, *P < 0.05 as determined by ANOVA.

We observed that the percentage of CD3+CD4+ but not CD3+CD8+ T cells was decreased in LN under the IL-2 hydrogel treatment (**Figure 3A,B**). In LN, the three times weekly 25,000 IU IL-2 and the once weekly 25,000 IU IL-2 hydrogel injection significantly reduced CD3+CD8+ T cell numbers (**Figure 3B**). The percentage of CD3+CD4+FoxP3+ Treg was significantly increased in the setting of the IL-2 hydrogel and the three times weekly 25000 IU IL-2 treatment in mesenteric LN (**Figure 3C)**. The fact that these changes were not observed in the spleen (**Figure 3D-F**) suggests that the effects of the hydrogel IL-2 release may be local. The Foxp3 MFI was unchanged in LN and spleen (**Supplemental Figure 3A,B**). In the setting of IL-2/IL-2 antibody complexes CD3+CD8+ T cells were reduced and Treg were increased (**Supplemental Figure 4**). We also explored the impact of delivering IL-2/HI complexes in the absence of a hydrogel but found that this yielded results no different from IL-2 alone (data not shown) suggesting that a stable, sustained release platform is needed.

Together, these data indicate that hydrogel-mediated IL-2 delivery increases Treg *in vivo*. Importantly, the IL-2 hydrogel once weekly matched the expansion of Treg compared to IL-2 alone given three times a week.

### IL-2 delivery in the context of a hydrogel reduces diabetes onset in NOD mice

In parallel to these effects on Treg expansion, we also asked whether these treatment regimens could reverse or prevent autoimmune diabetes in NOD mice.

Using an established protocol (22), we first asked whether 5 daily injections of 25,000 IU IL-2 or one single injection of a 25,000 IU IL-2 hydrogel could reverse new-onset diabetes in NOD mice. In our hands, no treatments reversed new-onset diabetes (data not shown).

We next asked whether a modified version of this established regimen could prevent autoimmune diabetes. NOD mice were injected once weekly with either 25,000 IU IL-2 or 25,000 IU IL-2 delivered in a hydrogel. This treatment was administered to six-week-old mice for 15 weeks. A Kaplan-Meyer curve shows % non-diabetic mice in the different treatment groups over time (**Figure 4A**).

**Figure 4.**
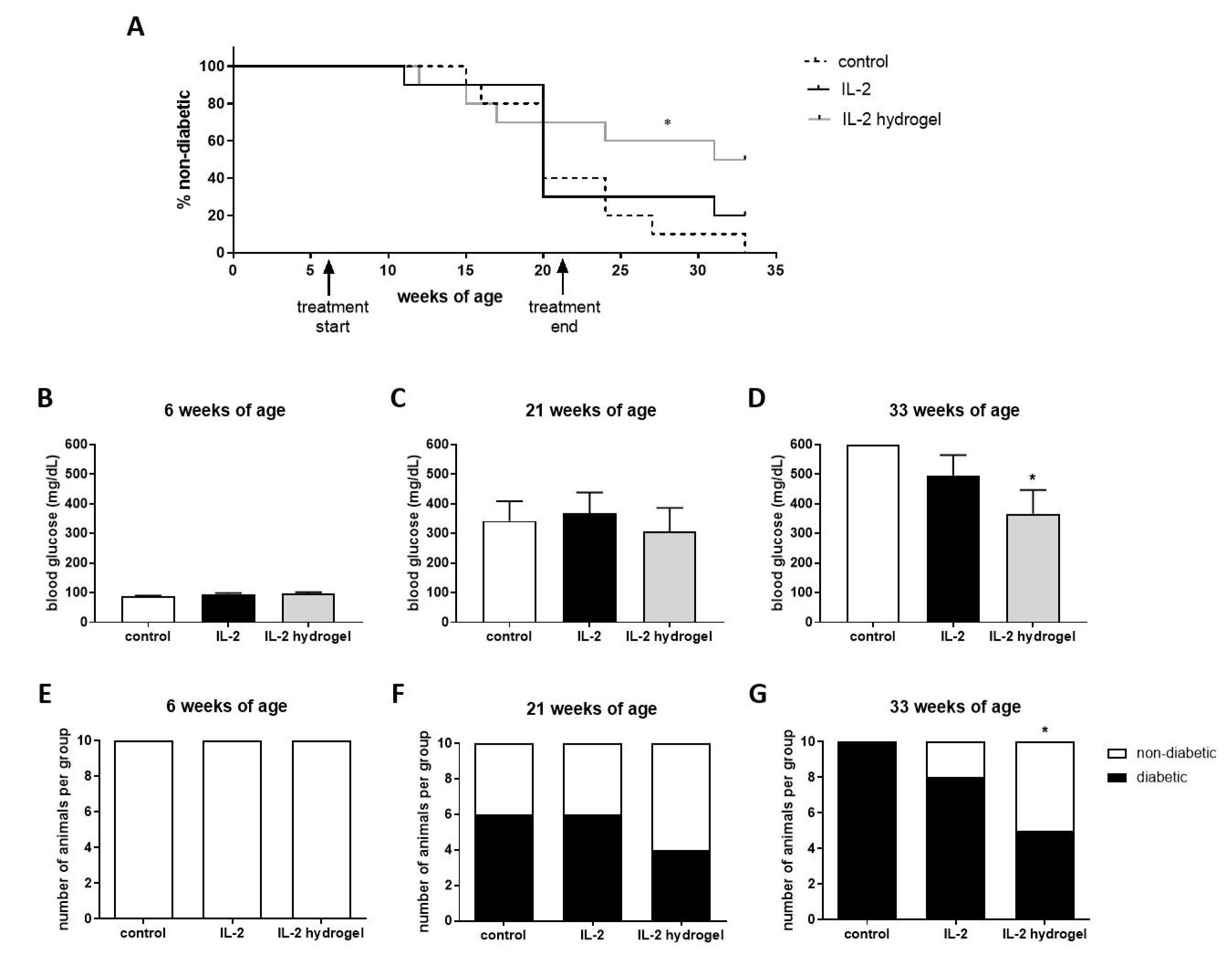
IL-2 delivery in the context of a hydrogel significantly reduces diabetes onset in NOD mice. **A.** Starting at 6 weeks of age, NOD mice received either 25,000 IU IL-2 once weekly, or the same amount of IL-2 delivered in the context of a once a week hydrogel injection. PBS injections served as negative controls. Mice were then monitored weekly for diabetes onset. This treatment was administered to mice from 6-21 weeks of age and mice were subsequently monitored until 33 weeks of age. **B-D.** Blood glucose at various ages, at 6 weeks when treatment started (**B**), at 21 weeks when the 15 week treatment period ended (**C**) and at 33 weeks of age, 3 month after treatment ended (**D**). **E-G**. Same data as in **B-D**, now expressed as number of diabetic mice, for 6 weeks of age (**E**), 21 weeks of age (**F**) and 33 weeks of age (**G**). N = 10 mice per group. Data represent mean +/- SEM, *P < 0.05 as determined by ANOVA.

After 15 weeks of treatment at 21 weeks of age, 6 out of 10 mice in the control and IL-2 treatment groups were diabetic (**Figure 4C, F**). In the IL-2 hydrogel treatment group 4 out of 10 mice were diabetic (**Figure 4C, F**). Three months after the end of the treatment, at 33 weeks of age, all mice of the control group, 8 out of 10 in the IL-2 group, and only 5 out of 10 in the IL-2 hydrogel group were diabetic (**Figure 4G**). At 33 weeks of age, the blood glucose values averaged >600 mg/dL for the control animals, ∼500 mg/dL for the IL-2 treatment group and ∼370 mg/dL for the IL-2 hydrogel group (**Figure 4D**). These data suggest that sustained release of IL-2 may enhance the long-term efficacy of treatment in this model.

## DISCUSSION

We report in a mouse model of T1D that an injectable hydrogel can be used to deliver IL-2 in a sustained manner.

This approach may enhance the effects of Treg expansion protocols as HI appears to potentiate the impact of IL-2 on Treg induction. A once-weekly IL-2 hydrogel injection was as effective as three-times-weekly IL-2 injections at inducing Treg.

Furthermore, these data suggest that it may be possible to reduce the onset of autoimmune diabetes using similar IL-2-releasing hydrogel preparations. We found that once-weekly IL-2 hydrogel injections prevented autoimmune diabetes in NOD mice by 50%. Moreover, we did not observe extensive CD8+ T cell activation. These data suggest that hydrogel-mediated controlled release of IL-2 may be a useful tool for IL-2 delivery.

This hydrogel IL-2 release material may have potential in treating other autoimmune disorders that are responsive to IL-2 (23), particularly perhaps autoimmune diseases of the skin. However, the safety and tolerability of these compounds in the context of IL-2 will need to be examined as well to avoid eosinophilic responses and other untoward effects (24,25). Moreover, it would be important to evaluate the performance of these hydrogel materials over a range of IL-2 concentrations and formulations.

We conclude that hydrogel-mediated IL-2 delivery may be a useful approach for delivering IL-2 in a sustained manner for use in treating autoimmunity.

## ACKNOWLEDGEMENTS

This work was supported in part by the Deutsche Forschungsgemeinschaft (DFG) NA 965/2-1 to NN and KA 3441/1-1 to GK; and National Institutes of Health grants R01 DK096087-01, R01 HL113294-01A1, and U01 AI101984 to PLB. KY was supported by the Swiss National Science Foundation early postdoctoral mobility grant and Child Health Research Institute and the Stanford NIH-NCATS-CTSA (grant no. UL1 TR001085). This work was also supported by grants from the JDRF 3-PDF-2014-224-A-N to NN and 1-SRA-2018-518-S-B Innovation Award to PLB and by grants from the Harrington Institute, Stanford SPARK, the Stanford Child Health Research Institute all to PLB.

## AUTHOR CONTRIBUTIONS

Conceptualization, NN, PLB; Investigation, NN, GK, MJK; Visualization, NN, HFK, SMR, PLB; Data/sample requisition, NN, GK, MJK, HFK, SMR, KY, JY; Writing, NN, MJK, PLB; Funding, NN, GK, KY, PLB; Supervision, NN, SCH, SAL, AP, PLB. All authors reviewed the results and approved the final version of the manuscript.

## ABBREVIATIONS

HA: hyaluronan
HI: heparin
IL-2: Interleukin 2
NOD: non-obese diabetic
T1D: Type 1 Diabetes

