## Supplemental Figures for "Sustained Release of IL-2 Using an Injectable Hydrogel Prevents Autoimmune Diabetes"

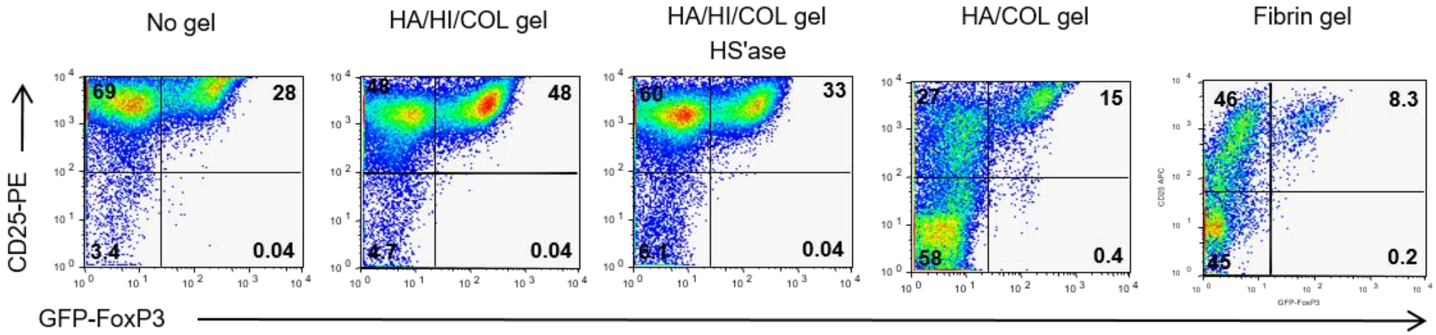


**Supplemental Figure 1. Heparin (HI) containing hydrogels potentiate Treg induction.** Representative flow plots of % Live FoxP3+ cells measured in different gel conditions. From the left to the right, no gel, gel containing HA/HI/COL, gel containing HA/HI/COL HS'ase digested, gel containing HA/COL and a fibrin gel.

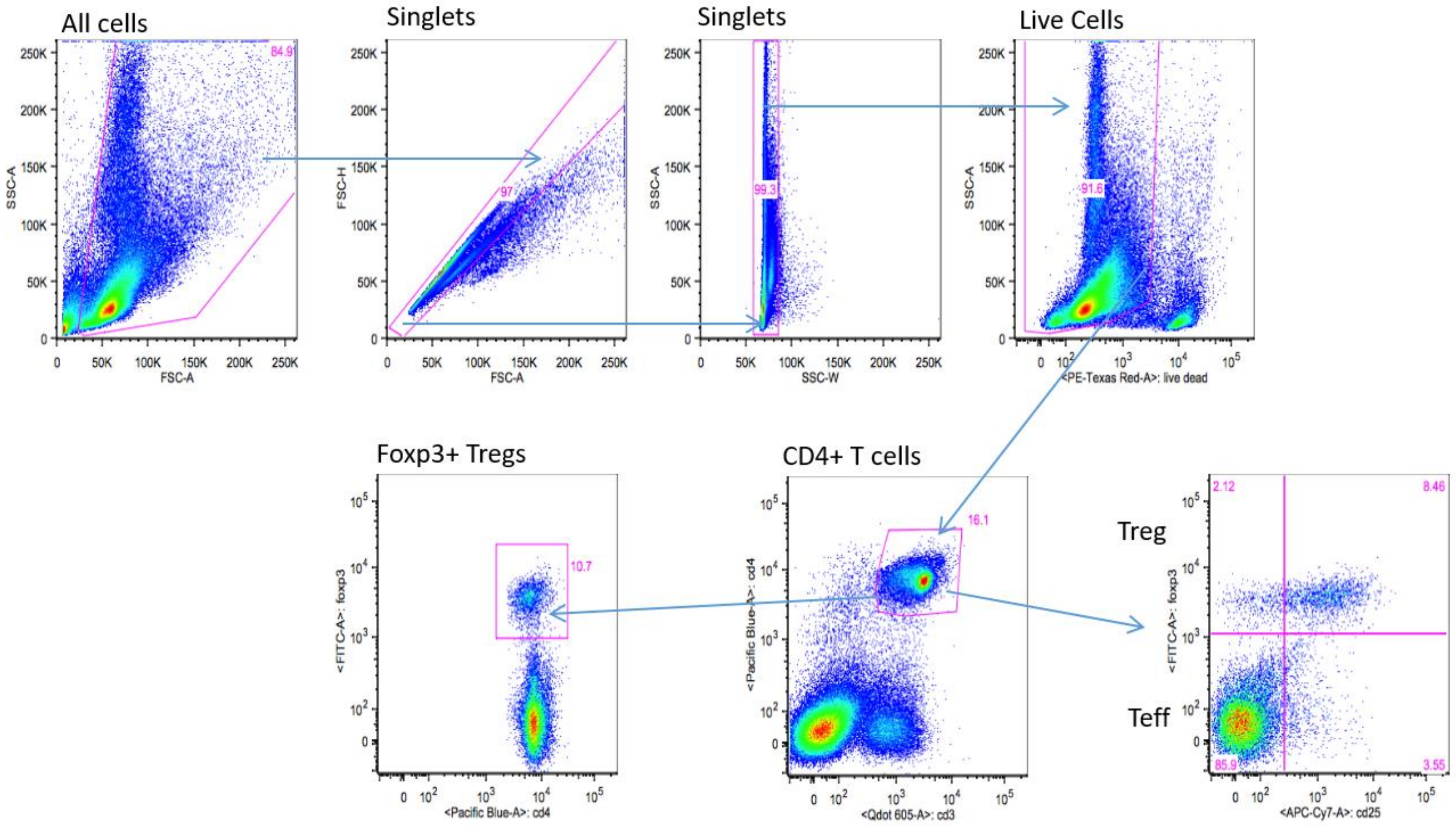


**Supplemental Figure 2. Gating scheme for Figure 3.** Gating scheme for the quantification of Foxp3+ regulatory T cell numbers in mice. Lymphocyte and splenocyte preparations were stained according to a 6-color staining panel and analyzed by flow cytometry. Live lymphocytes and splenocytes were successively gated to identify Foxp3+ regulatory T cells amongst CD3+/CD4+ cells.

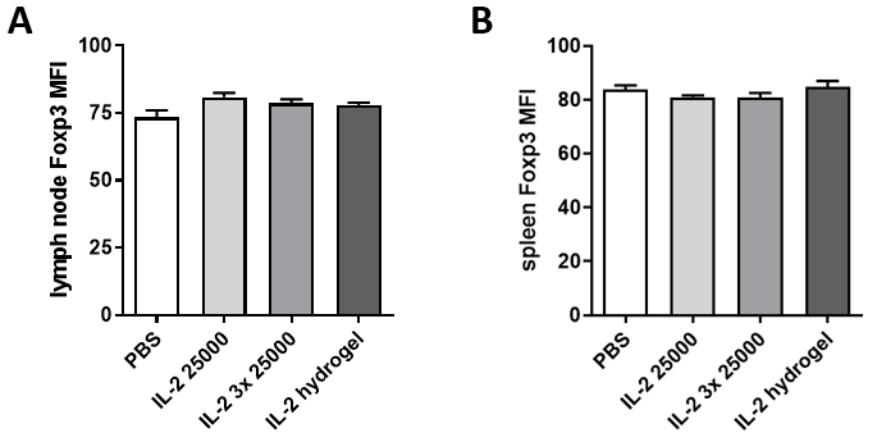


**Supplemental Figure 3. Hydrogel-mediated IL-2 delivery is not associated with an increase in FoxP3+ Treg MFI.** 6-week-old NOD mice were treated with either 25,000 IU IL-2 once weekly or three times a week, the same amount of IL-2 delivered in the context of a single hydrogel injection, or PBS injections as a negative control. After one month, mice were euthanized and both mesenteric lymph nodes (**A**) and spleens (**B**) were collected and populations of lymphocytes were assessed by flow cytometry. In particular, the CD3+CD4+FoxP3+ Treg MFI (**A, B**), Data are representative of 2 independent experiments. N = 10 mice per group. Data represent mean +/- SEM.

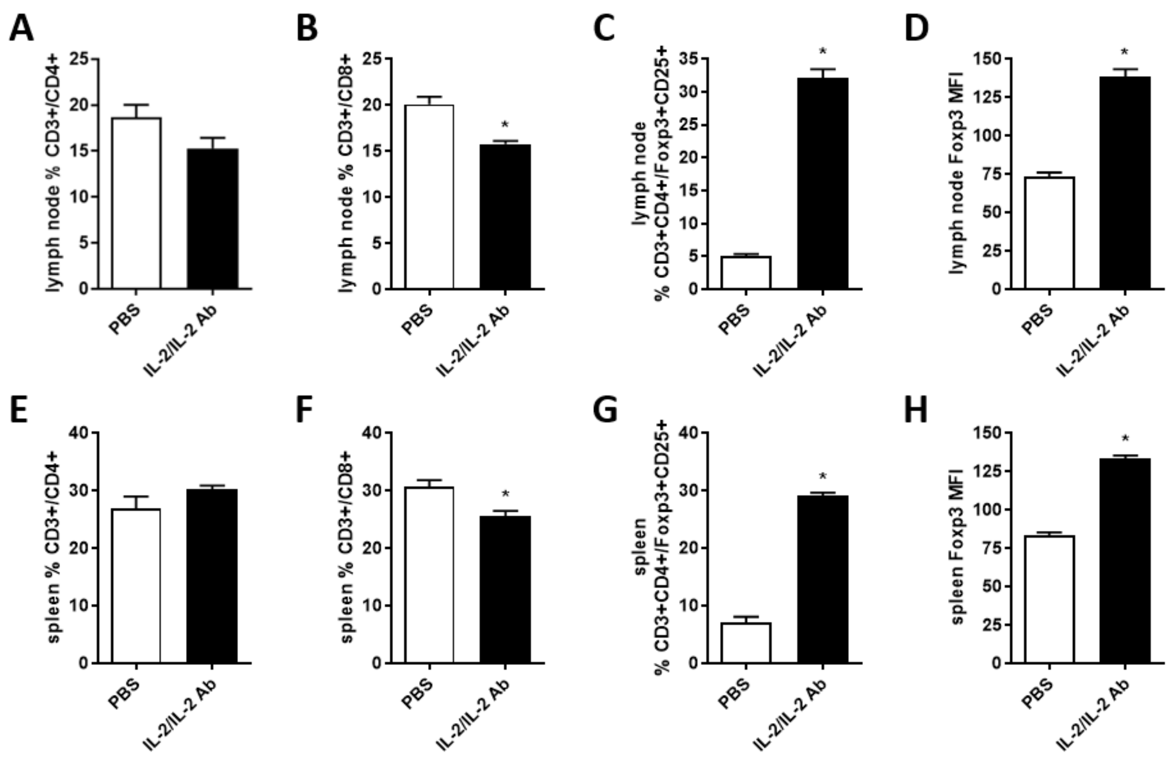


**Supplemental Figure 4. IL-2 control for IL-2 hydrogels.** 6-week-old NOD mice were treated with PBS injections as a negative control or IL-2/IL-2 Ab as positive control. After one month, mice were euthanized and both mesenteric lymph nodes (**A-D**) and spleens (**E-H**) were collected and populations of lymphocytes were assessed by flow cytometry. In particular, the percentage of CD3+CD4+ T-cells (**A,E**), the percentage of CD3+CD8+ T-cells (**B,F**) and CD3+CD4+FoxP3+ Treg (**C,G**) and Foxp3+ MFI (**D,H**). Data are representative of 2 independent experiments. N = 10 mice per group. Data represent mean +/- SEM, *P < 0.05 as determined by ANOVA.
